# Peptide-based quorum sensing systems in *Paenibacillus polymyxa*

**DOI:** 10.1101/767517

**Authors:** Maya Voichek, Sandra Maaß, Tobias Kroniger, Dörte Becher, Rotem Sorek

**Affiliations:** Department of Molecular Genetics, Weizmann Institute of Science, Rehovot 76100, Israel; Institute of Microbiology, Department of Microbial Proteomics; Center for Functional Genomics of Microbes, University of Greifswald, Germany

## Abstract

*Paenibacillus polymyxa* is an agriculturally important plant growth-promoting rhizobacterium. Many *Paenibacillus* species are known to be engaged in complex bacteria-bacteria and bacteria-host interactions, which in other species were shown to necessitate quorum sensing communication. However, to date no quorum sensing systems have been described in *Paenibacillus*. Here we show that the type strain *P. polymyxa* ATCC 842 encodes at least 16 peptide-based communication systems. Each of these systems is comprised of a pro-peptide that is secreted to the growth medium and processed to generate a mature short peptide. Each peptide has a cognate intracellular receptor of the RRNPP family, and we show that external addition of *P. polymyxa* communication peptides leads to reprogramming of the transcriptional response. We found that these quorum sensing systems are conserved across hundreds of species belonging to the *Paenibacillaceae* family, with some species encoding more than 25 different peptide-receptor pairs, representing a record number of quorum sensing systems encoded in a single genome.

## Introduction

*Paenibacillaceae* is a diverse family of bacteria, many of which are important in agricultural and clinical settings. These include *Paenibacillus larvae*, a pathogen causing the lethal American Foulbrood disease in honeybees^1^, as well as *P. dendritiformis* and *P. vortex*, which are used as models for complex colony pattern formation^2,3^. Arguably the best studied member of *Paenibacillaceae* is *P. polymyxa*, known for its plant-growth promoting traits^4–6^. *P. polymyxa* produces a plethora of beneficial compounds, including the antibiotic polymyxin^7^ and the phytohormone indole-3-acetic acid (IAA)^8^, and was shown to protect multiple plant species against pathogens (reviewed in ^9^). Due to its beneficial traits, *P. polymyxa* is commercially used as a soil supplement to improve crop growth^10,11^.

Many of the characteristic features associated with the above-mentioned behaviors – production and secretion of antimicrobials, expression of virulence factors, and forming complex colony structures – often require some form of inter-cellular communication such as quorum sensing^12^. Bacteria use quorum sensing to coordinate gene expression patterns on a population level. For example, the quorum sensing network of *Pseudomonas aeruginosa* controls the production of secreted toxins as well as the production of rhamnolipids, which are important for its biofilm architecture^13,14^. Quorum sensing regulates the production of bacteriocin antimicrobials in *Lactobacilli*^15^ and *Streptococci*^16,17^; and in *Bacillus subtilis*, quorum sensing systems play a major role in sporulation, biofilm formation and genetic competence^18^. Although *Paenibacillus* bacteria are engaged in similar behaviors, no quorum sensing was reported in any *Paenibacillus* species to date.

Gram-positive bacteria frequently use short peptides, termed autoinducer peptides, as their quorum sensing agents^12^. A widespread group of such peptide-based communication systems involves intracellular peptide sensors (receptors) of the RRNPP family, together with their cognate peptides^19^. Peptides of these quorum sensing systems are secreted from the cell as pro-peptides and are further processed by extracellular proteases^20^ to produce a mature short peptide, usually 5-10 amino acids (aa) long^19,21^. The mature peptides are re-internalized into the cells via the oligopeptide permease (OPP) transporter^22,23^, and bind their intracellular receptors^24^. These receptors act either as transcription factors or as phosphatases, and peptide binding activates or represses their function eventually leading to modulation of the bacterial transcriptional program. For example, in *B. subtilis* the Rap receptors function as phosphatases that regulate the major transcription factor Spo0A; upon binding to their cognate Phr peptides, the phosphatase activity of Rap receptors is inhibited, leading to activation of Spo0A that alters the transcriptional program^25,26^. In *B. thuringiensis* the PlcR receptor is a transcription factor that becomes activated when bound to its cognate PapR peptide, leading to expression of virulence factors that facilitate infection of its insect larval host^27^. Such peptide-based communication was recently shown also to control the lysis-lysogeny decision of phages infecting many species of *Bacillus*^28,29^.

Here we report the identification of a large group of peptide-based quorum sensing systems in *P. polymyxa*. We found that pro-peptides of these systems are secreted and further processed, and that the mature peptides modulate the transcriptional program of *P. polymyxa*. We further show that these quorum sensing systems are conserved throughout the *Paenibacillaceae* family of bacteria, with some bacteria encoding over 25 different peptide-receptor pairs in a single genome.

## Results

### Identification of RRNPP-like peptide-receptor pairs in P. polymyxa ATCC 842

All members of the RRNPP family of intracellular peptide receptors contain a C-terminal tetratricopeptide repeat (TPR) or TPR-like domain, which forms the peptide-binding pocket in the receptor protein^19,30^. To search for possible such quorum sensing systems in *P. polymyxa* we first scanned all annotated protein-coding genes of the type strain *P. polymyxa* ATCC 842^31^ for TPR domains using TPRpred^32^ (Methods). As TPR domains are found in diverse protein types, we manually analyzed the genomic vicinity of the identified TPR-containing proteins to search for a peptide-encoding gene characteristic of quorum sensing systems. We found two TPR domain genes that were immediately followed by a short open reading frame with a predicted N-terminal hydrophobic helix, typical of a signal peptide that targets the pro-peptide for secretion^33^ (Fig. 1a). Two-gene operons that encode the receptor followed by its cognate peptide are a hallmark of many known members of the RRNPP family of quorum sensing systems^19^.

**Fig. 1:**
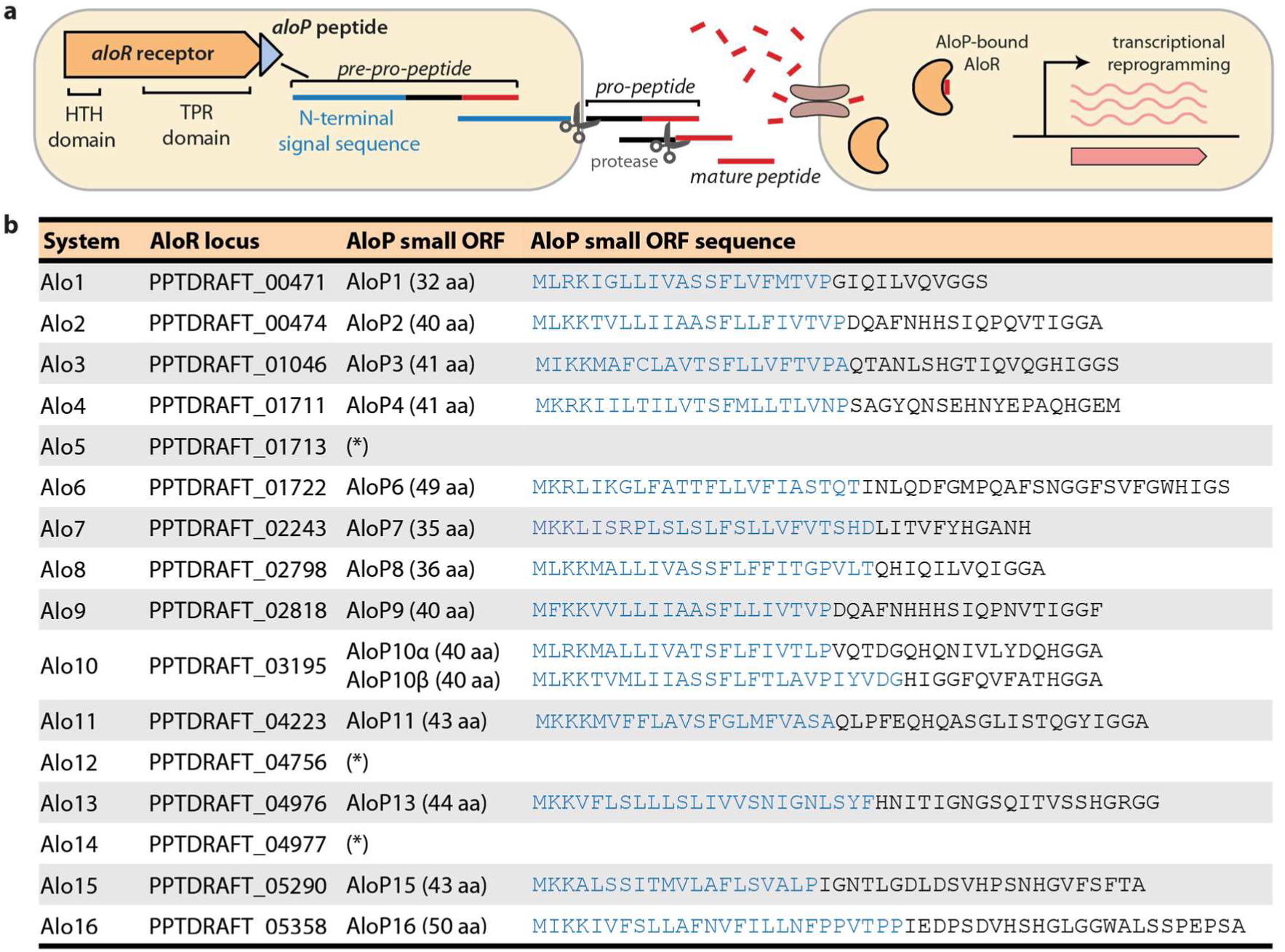
Alo systems identified in *P. polymyxa*. (a) Schematic representation of *aloR*-*aloP* operon organization and putative peptide processing. HTH, helix-turn-helix; TPR, tetratricopeptide repeat. (b) Alo loci identified in *P. polymyxa* ATCC 842. For the *aloR* receptor genes, the locus tag in the IMG database^34^ is specified. Blue sequence in AloP peptides represents the predicted N-terminal signal sequence for secretion, as predicted by Phobius^35^ (Methods). Asterisks (*) mark cases in which the *aloP* gene was absent or found as a degenerate sequence.

We then used BLAST to search for additional homologs of the two putative TPR-domain receptors in the *P. polymyxa* ATCC 842 genome (Methods). This search revealed 14 additional homologous genes that were not initially recognized by TPRpred, in most cases because their C-terminal TPR domains were divergent and did not pass the default threshold of the TPRpred software. We found that all but three of the homologs had a short ORF (32-50 aa) encoded immediately downstream to them. We denote these putative communication systems as Alo (<u>A</u>utoinducer peptide <u>Lo</u>cus), with predicted Alo receptor genes designated *aloR* and the cognate peptide-encoded gene *aloP*. We number Alo loci in the *P. polymyxa* ATCC 842 genome from *alo1* to *alo16* (Fig. 1b).

### Alo systems are ubiquitous in P. polymyxa strains

Examining the genomes of 13 additional *P. polymyxa* strains using sequence homology searches revealed that Alo systems are conserved in the *P. polymyxa* lineage (Fig. 2a; Methods). Some of the Alo receptors (e.g., AloR3, AloR4, AloR7, AloR13, AloR15, and AloR16) are highly conserved, and appeared in all *P. polymyxa* strains examined. Others appeared variably and were absent from some genomes (Fig. 2b). We also found six additional peptide-receptor pairs that were not present in *P. polymyxa* ATCC 842 but were detectable in other strains; we denote these Alo17-22 (Fig. 2). Similar observations regarding “core” and “accessory” quorum sensing systems were reported for peptide-based systems in other bacteria: for example, in the case of *B. subtilis* Rap/Phr genes, RapA/PhrA and RapC/PhrC are considered “core” systems that are conserved among *B. subtilis* species, while other systems, including RapE/PhrE, RapI/PhrI and RapK/PhrK, occur variably and are often located within mobile genetic elements^36^.

**Fig. 2:**
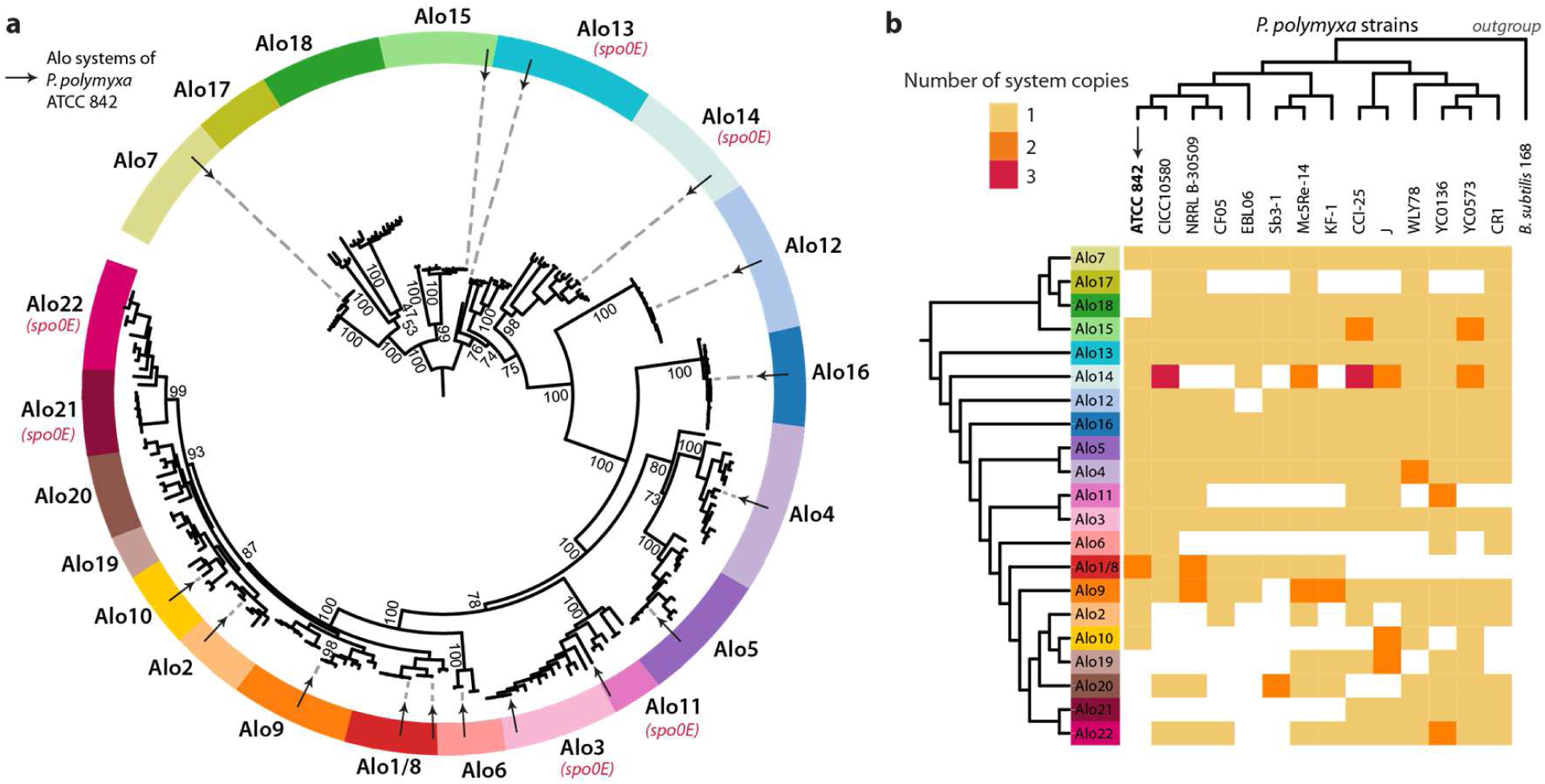
Phylogenetic distribution of Alo systems among *P. polymyxa* strains. (a) Phylogenetic tree of 234 AloR homologs found in 14 *P. polymyxa* strains. Colored segments in the tree circumference define clades of AloR proteins corresponding to the systems found in *P. polymyxa* ATCC 842 (labeled as arrows). Systems immediately followed by a Spo0E-like protein are indicated. Tree was constructed using IQ-Tree^38–40^ with 1,000 iterations and ultrafast bootstrap support values are presented for major branches. Tree visualization was done with iTOL^41^. Alo1/8 refers to a system duplicated in *P. polymyxa* ATCC 842 (where it is represented as Alo1 and Alo8). (b) Frequency and distribution of Alo systems in *P. polymyxa* strains. The number of copies per each Alo system found within *P. polymyxa* strains is shown using colored squares. The absence of an Alo system in a specific strain is marked by a white square. Phylogenetic tree of the Alo systems shown on the vertical axis is derived from the tree in panel (a). Phylogenetic tree of the strains shown on the horizontal axis is based on the GyrA protein, with GyrA from *B. subtilis* 168 as an outgroup.

In the vast majority of cases, *aloR* genes in the various *P. polymyxa* strains were found just upstream to a peptide-encoding *aloP* gene with an N-terminal hydrophobic helix predicted to target them for secretion by the Sec system^26^ (Supplementary Table S1). Exceptions were *aloR18*, for which we could not find a cognate downstream short ORF, and *aloR5* and *aloR14* in which degenerate peptide sequences were observed. In the Alo12 clade we could identify a conserved short ORF in the majority of *P. polymyxa* strains (excluding the type strain ATCC 842), yet it lacked the characteristic hydrophobic N-terminal signal sequence marking it for secretion. Orphan receptors, which lack a cognate peptide, have also been observed for RRNPP quorum sensing systems in other bacteria^36,37^.

To understand how common Alo systems are in bacteria, we performed an exhaustive profile-based search for homologs of AloR proteins in >38,000 bacterial and archaeal genomes (Methods). This search yielded 1,050 homologs in 149 bacteria, essentially all of which belong to the *Paenibacillaceae* family (Supplementary Table S2). These included species of *Paenibacillus, Brevibacillus, Saccharibacillus, Fontibacillus* and *Gorillibacterium*. The largest number of Alo receptors found in a single organism was in *Paenibacillus terrae* NRRL B-30644, in which 40 such receptors were detected (with cognate peptide-encoding genes detected immediately downstream to 27 of them, Supplementary Table S3). These results suggest that the Alo system is a widespread quorum sensing system in this large family of bacteria, and imply that *Paenibacilli* are among the most “communicative” bacteria studied to date.

### Mass-spectrometry analysis detects secretion and processing of pro-peptides

Peptides belonging to members of the RRNPP family of quorum sensing systems are known to be secreted into the growth medium as pro-peptides. Following removal of their N-terminal signal sequence, the peptides are further processed by extracellular proteases to generate the mature communication peptide^26,42^ (Fig. 1a). To determine whether Alo peptides are similarly secreted, we analyzed growth media taken from *P. polymyxa* cultures using mass-spectrometry (MS). For this, we grew *P. polymyxa* ATCC 842 in defined media, removed the bacteria by centrifuging, and took the supernatant (presumably containing the secreted peptides) for MS analysis. Notably, MS was performed without subjecting the peptides to trypsin treatment, to allow detection of peptides in their natural form. We were able to reproducibly detect various fragments of nine out of the 14 Alo peptides predicted in *P. polymyxa* ATCC 842 (Fig. 3 and Supplementary Table S4). In all cases, the fragments included the C-terminus of the Alo peptide but not the N-terminus, confirming that the N-terminus is cleaved during the pro-peptide secretion, presumably by a signal peptidase associated with the Sec system^43^. As a control, we repeated the same procedure on cultures of *B. subtilis*, where the processing patterns of Phr peptides are known^26^. Indeed, the *B. subtilis* Phr peptides were also detectable in the MS analysis, with the same patterns of N-terminus removal (Fig 3A and Supplementary Table S4). These results show that Alo peptides are secreted to the medium akin to other cases of RRNPP quorum sensing systems.

**Fig. 3:**
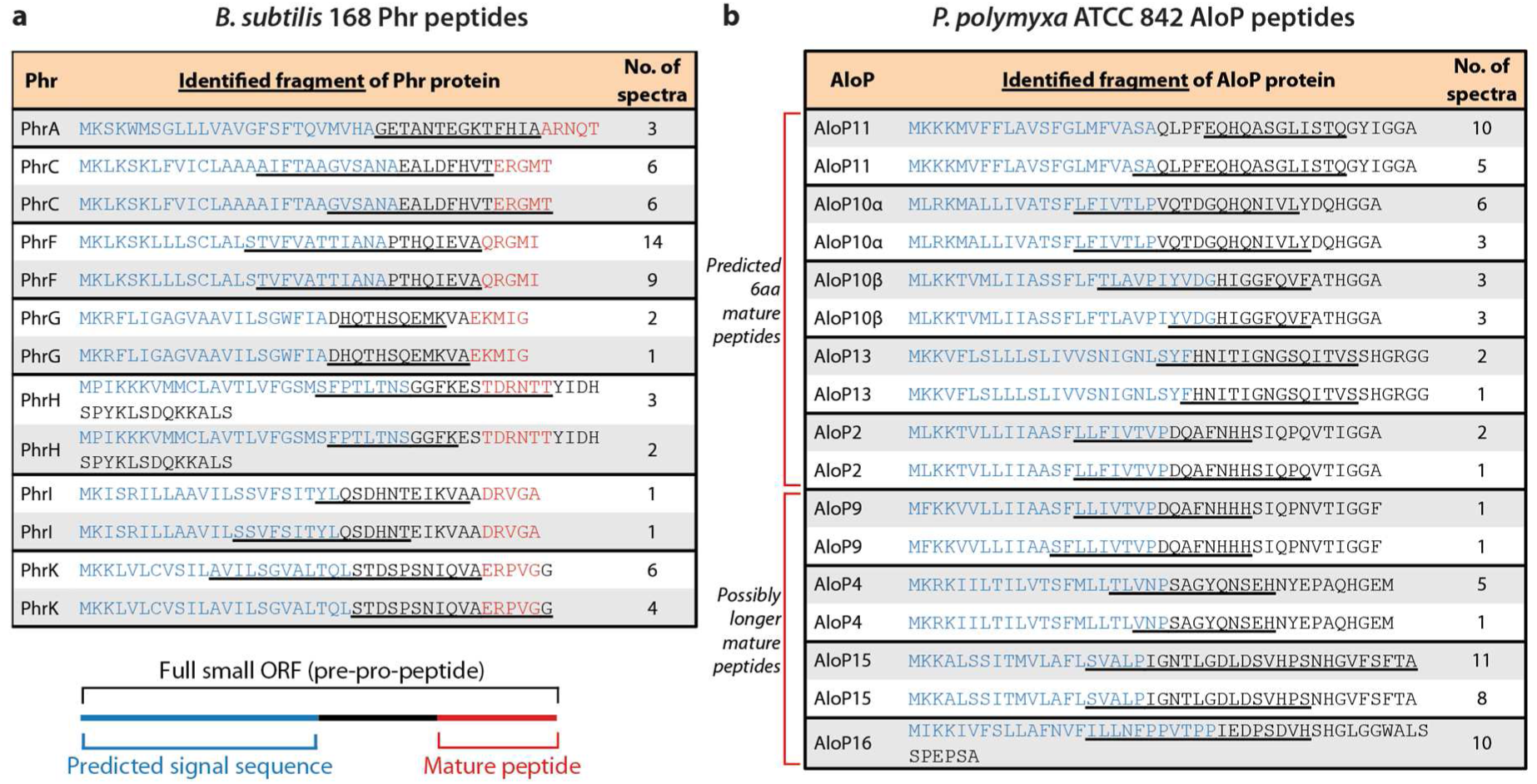
Mass spectrometry-based identification of secreted communication peptides. (a) Phr peptide fragments detected in the growth media of *B. subtilis* 168. Peptide fragments identified by MS are underlined, with the number of spectra detected indicated. Known mature communication peptides are in red^26,45^. Blue indicates the predicted N-terminal signal sequence for secretion^26^. The two most abundantly identified fragments are presented for each protein. (b) AloP peptide fragments detected by MS. Same as (a) for AloP peptide fragments detected in the growth media of *P. polymyxa* ATCC 842. Blue indicates the N-terminal signal sequence for secretion as predicted by Phobius^35^.

Following N-terminus removal during the secretion process, *B. subtilis* Phr peptides are known to be processed further to yield the mature 5 aa peptide. The mature peptide is usually found at the extreme C-terminal end of the pro-peptide, such that it may be released by a single cleavage event (Fig. 3a). Such short peptides are challenging to identify using current MS pipelines^44^, and indeed our MS analysis of *B. subtilis* conditioned media did not detect the mature 5 aa Phr peptides but rather revealed the Phr pro-peptides with the known mature peptide missing (Fig. 3a). The absence of a short peptide from the C-terminus of the pro-peptide is therefore a clear signature of the processing event that releases the mature peptide. We were able to detect similar processing events in the *P. polymyxa* peptides as well (Fig. 3b). These patterns strongly suggest that some of the AloP proteins generate 6 aa communication peptides, while others generate presumably longer mature peptides (Fig. 3b).

### Peptide-mediated modulation of the bacterial transcriptional program

All Alo receptors contain a helix-turn-helix (HTH) DNA-binding domain at their N-terminus (Fig. 1a), suggesting that they may function as transcriptional regulators similar to many members of the RRNPP peptide receptors family^19,24^. These regulators become activated or repressed once bound by their cognate peptides, leading to alteration of the transcriptional program in response to the quorum sensing peptides. To examine whether Alo peptides affect the transcriptional program of *P. polymyxa*, we selected Alo13, one of the systems conserved across all *P. polymyxa* strains, for further experimental investigation. The sequence of the mature AloP13 peptide, as predicted by the MS analysis, is SHGRGG (Fig. 3b). Aiming to measure the immediate transcriptional response to the peptide, we incubated early log-phase-growing *P. polymyxa* cells with 5µM of synthetic SHGRGG peptide for 10 minutes. RNA-seq analysis of cells harvested after this short incubation with the peptide identified a set of 10 genes whose expression became significantly reduced when the peptide was added (Fig. 4). This transcriptional response was not observed when a scrambled version of the AloP13 pro-peptide (encoding the same amino acids as the pro-peptide but in a scrambled order) was added to the medium, indicating that the transcriptional response is specific to the sequence of AloP13 (Table S5).

**Fig. 4:**
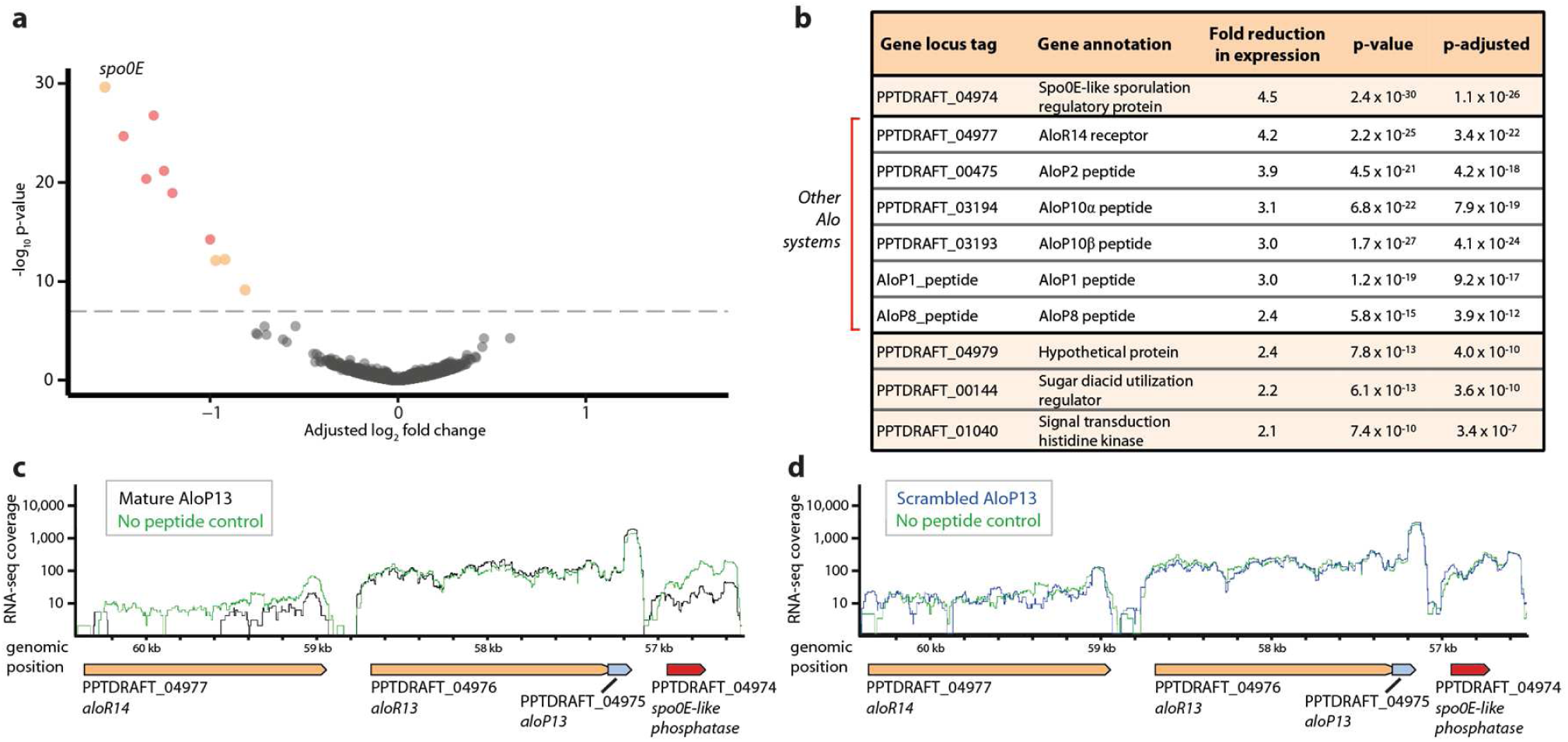
The mature AloP13 peptide elicits an immediate transcriptional response. (a) Volcano plot depicting *P. polymyxa* gene expression after incubation with 5µM of the peptide SHGRGG for 10 minutes, as compared to control conditions in which no peptide was added. X-axis, log_2_ fold expression change (adjusted using the lfcShrink function in the DESeq2 R package^48^, Methods). Y-axis, −log_10_ p-value. Average of 3 independent replicates; each dot represents a single gene. Dots appearing in colors are genes that passed the threshold for statistical significance of differential expression (Methods). Red dots correspond to genes encoded by Alo systems. (b) Differentially expressed genes. Fold change represents the average of the 3 independent replicates. P-adjusted is the p-value after correction for multiple hypothesis testing (Methods). (c) RNA-seq coverage of the Alo13-Alo14 locus (Scaffold: PPTDRAFT_AFOX01000049_1.49), in control conditions (green) or 10 minutes after addition of 5 µM of the SHGRGG peptide (black). RNA-seq coverage is in log scale and was normalized by the number of uniquely mapped reads in each condition. Representative of 3 independent replicates. (d) RNA-seq coverage of the Alo13-Alo14 locus in control conditions (green) or 10 minutes after addition of 5µM of the scrambled AloP13 pro-peptide VSGTRHGSFHSGIGNSGIYIQNT (blue). RNA-seq coverage is in log scale and normalized as in panel (c). Representative of 3 independent replicates. Data for the control sample is the same as in panel (c).

The most significant reduction in expression was in a gene annotated as Spo0E-like sporulation regulatory protein, whose expression was reduced by 4.5 fold on average in response to the peptide (3.7, 6, and 4 fold reduction in the 3 independent replicates of the experiment) (Fig. 4b). In *B. subtilis*, Spo0E functions as a phosphatase that regulates Spo0A, the master transcription factor for sporulation, thus inhibiting the initiation of the sporulation pathway^46^. Interestingly, the *spo0E*-like gene is encoded directly downstream to the Alo13 locus in *P. polymyxa* (Fig. 4c-d). These results imply that the immediate effect of the AloP13 quorum sensing peptide involves down regulation of the *spo0E*-like gene, which likely leads to a further regulatory cascade at later time points. We note that *spo0E*-like genes occur immediately downstream of 6 Alo systems in *P. polymyxa* (Fig. 2a) and downstream to 12 of the *P. terrae* Alo systems. This conserved genomic organization suggests that many Alo systems may exert their effect on the cell by modulating the expression of Spo0E-like regulatory phosphatase proteins.

Most of the additional genes whose expression became reduced following 10 minutes’ exposure to the AloP13 mature peptide were those encoding other AloP peptides, including *aloP1, aloP2, aloP8, aloP10α* and *aloP10β*, as well as the gene encoding the receptor *aloR14* that is encoded adjacent to the *aloR13* gene (Fig. 4). These results suggest that as in other bacteria^37,47^, quorum sensing systems can cross-regulate other such systems in *P. polymyxa*. Combined, these results show that the AloP13 mature peptide affects the transcription of specific target genes, and verifies Alo systems as quorum sensing systems in *P. polymyxa*.

### The AloP13 pro-peptide induces bacterial clumping

We noticed that addition of the AloP13 pro-peptide (sequence SYFHNITIGNGSQITVSSHGRGG) to a liquid growth medium resulted in rapid clumping of the bacteria into macroscopic particles (Fig. 5; Supplementary Movie S1). The clumping phenotype repeated when using an AloP13 pro-peptide synthesized by two different vendors. Clumping was not observed when using the synthetic “scrambled” peptide encoding the same amino acids as the AloP13 pro-peptide but in a scrambled order, suggesting that clumping is specifically induced by the AloP13 pro-peptide (Fig. 5). Clumping was previously described to be mediated by quorum sensing in other bacteria, where it was associated with induction of conjugation^49^, competence^50^ and virulence^51^. Surprisingly, addition of the mature 6 aa peptide SHGRGG to the medium did not result in observable clumping, and, conversely, addition of an AloP13 pro-peptide that lacks the last 6 aa did induce clumping (Fig. 5b). These results suggest that the AloP13 pro-peptide carries out two separate functions – one is associated with the mature SHGRGG peptide that probably affects transcription by entering the cell and binding its cognate receptor, and the other induces bacterial clumping in a mechanism yet to be explored.

**Fig. 5:**
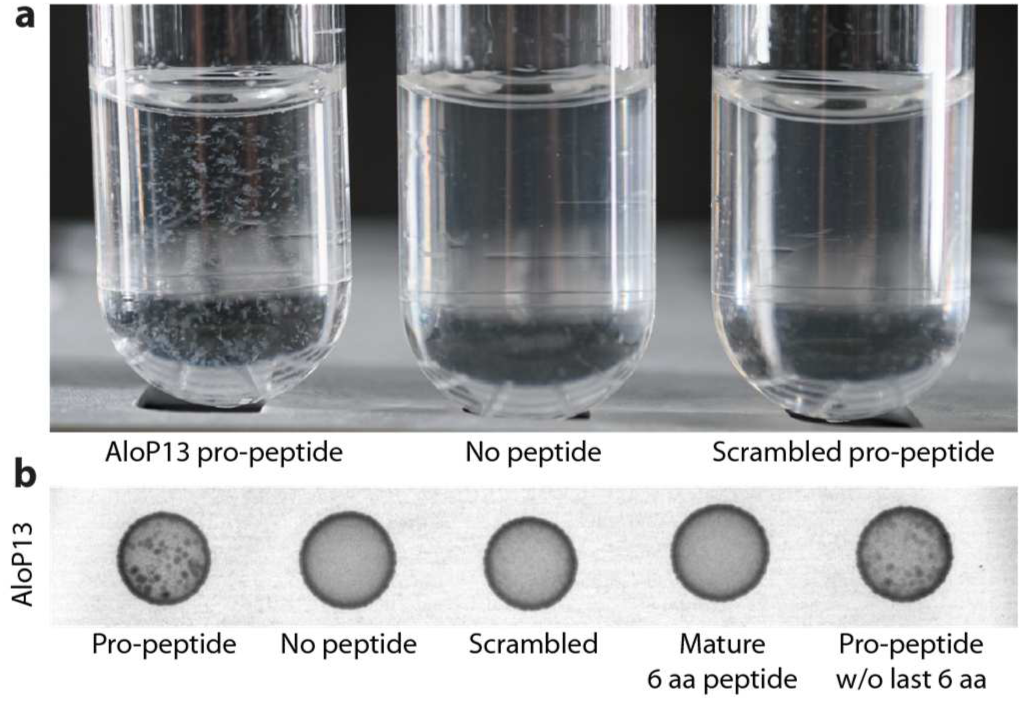
The pro-peptide of AloP13 causes cell clumping. (a) Cell clumping is observed in liquid culture of *P. polymyxa*, 30 minutes after addition of 5µM of the AloP13 pro-peptide SYFHNITIGNGSQITVSSHGRGG. Clumping is not observed in a control sample in which no peptide was added, nor in a sample incubated for 30 minutes with 5µM of a scrambled version of the AloP13 pro-peptide (VSGTRHGSFHSGIGNSGIYIQNT). (b) Cultures grown similarly to those in panel (a), as well as cultures incubated with the mature AloP13 (SHGRGG) and an AloP13 lacking the last 6 aa (SYFHNITIGNGSQITVS) were plated on LB-agar and grown overnight. Clumping is observed as denser areas in the culture. Image in inverted colors.

## Discussion

In this work we identified a large family of peptide-based quorum sensing systems employed by *Paenibacillaceae* bacteria, and characterized these systems in the type strain *P. polymyxa* ATCC 842. We found that the peptides are secreted to the growth medium and further processed to form mature communication peptides which lead to transcriptional reprogramming. The large number of such systems in a single bacterium (16 in *P. polymyxa* ATCC 842 and at least 27 in *P. terrae*) suggest that Paenibacilli may need an unusually large amount of signals to coordinate their complex social traits. Previous studies have predicted that *P. polymyxa* strains may also utilize AI-2 quorum sensing, based on the presence of components of the AI-2 pathway in its genome, although the production of AI-2 by *P. polymyxa* was not detected^52,53^. In addition, *P. polymyxa* ATCC 842 was found to encode an *agr*-like gene cassette that is homologous to another peptide-based communication system in staphylococci^54^. Together with our discovery of the Alo communication systems, to our knowledge, Paenibacilli have the largest number of quorum sensing systems reported in any bacteria to date.

In *B. subtilis* most Phr pro-peptides are processed by a single protease cleavage event, releasing the C-terminal mature peptide. However, in some cases there are two cleavage events (for example PhrH and PhrK), releasing an internal portion of the pro-peptide to form the mature communication peptide^26,45^. Our MS data of *B. subtilis* peptides show that these cases may not be detectable in our analyses, but instead may appear as longer mature peptides (see PhrH and PhrK in Fig. 3a). It is therefore conceivable that some of the AloP peptides in *P. polymyxa*, especially the ones for which the fragment detected in the MS data is followed by a sequence longer than 6 aa, are further processed to yield shorter mature peptides (for example AloP4, P9, P15, and P16 in Fig. 3b).

Our data suggest that AloP13 has two separate functions. First, its terminal 6 aa are processed and trigger transcriptional reprogramming similar to other peptides in the RRNPP family of quorum sensing systems. Second, another portion of its sequence (which does not include the last 6 aa) causes the cells to aggregate. There are other known cases of a communication peptide carrying two separate functions: for example, the bacteriocin nisin is known to act both as an antimicrobial peptide and as a signaling molecule that induces its own biosynthesis in a cell density-dependent fashion^55,56^. As for the observed phenotype, cell aggregation is known to be caused by the CSP communication peptide in *Streptococcus pneumoniae*, in which the peptide leads to cell competence for DNA uptake^50^. It is therefore possible that AloP13 regulates natural competence in *P. polymyxa* as well. If this is indeed the case, then using AloP peptides may facilitate DNA transformation and genetic engineering of *P. polymyxa*, which is currently considered a species challenging to genetically manipulate^57–59^.

We found that addition of AloP13 to growing cells results in immediate down-regulation of a set of genes, including those encoding the Spo0E-like phosphatase and additional Alo peptides. These immediate expression changes are expected to be translated into a signaling cascade that may involve alteration of the phosphorylation state of additional cellular components, ultimately giving rise to a functional phenotype. The phenotype controlled by AloP13, as well as those controlled by other AloP peptides, remain to be elucidated. Revealing the full network of functional phenotypes controlled by peptide communication in *P. polymyxa* may enable efficient harnessing of the plant-growth promoting traits of this organism for agricultural benefit in the future.

## Materials and Methods

### Identification of Alo systems in P. polymyxa ATCC 842

Amino acid (aa) sequences of all protein-coding genes of *Paenibacillus polymyxa* N.R. Smith 1105, ATCC 842 were downloaded from IMG^34^ (on 31/1/2017, IMG taxon ID: 2547132099, NCBI:txid 1036171) and scanned with TPRpred version 2.8^32^. Genes with TPRpred scores of >75% probability of having TPR/PPR/SLE domains were further inspected. The genome environments of these genes were manually searched for downstream annotated short ORFs with an N-terminal hydrophobic helix as predicted by the Phobius web server^35^. For two TPR-containing genes, such downstream short ORFs were identified. BlastP^60^ was then used to search for homologs of these two genes among all *P. polymyxa* ATCC 842 proteins using an E-value cutoff of 1e-5. Homologs shorter than 250 aa were discarded. The remaining 16 genes were numbered *aloR1*-*aloR16*, and their cognate *aloP* genes were searched as above. In cases no cognate *aloP* gene could be identified, the last 100 bases of the *aloR* gene together with the 200 bases immediately downstream of the *aloR* were searched for short ORFs (30-50 aa) partially overlapping the 3’ of the *aloR* gene using Expasy translate tool^61^. N-terminal hydrophobic helices of these AloP peptides were predicted using the Phobius web server^35^. Signal sequences were manually inferred in cases where the Phobius score was near the threshold based on homology to other AloP sequences whose Phobius score was above the threshold.

### Phylogenetic analysis of Alo systems

The aa sequences of the 16 AloR proteins identified in *P. polymyxa* ATCC 842 were used as a query for iterative homology search against 38,167 bacterial and archaeal genomes downloaded from the IMG database^34^ on October 2017, using the “search” option in the MMseqs2 package^62^ (release 6-f5a1c) with “--no-preload --max-seqs 1000 --num-iterations 3” parameters. The hits were filtered with an E-value cutoff of <1e-10. Homologs shorter than 250 aa or homologs found in scaffolds shorter than 2,500 nt were discarded.

For the analysis appearing in Fig. 2, the 234 AloR homologs found in 14 *P. polymyxa* strains were aligned using the MAFFT multiple sequence alignment server version 7 with default parameters^63^. The alignment was used to generate a maximum likelihood phylogenetic tree using IQ-TREE^38^ version 1.6.5 including ultrafast bootstrap analysis^39^ with 1000 alignments (‘‘-bb 1000’’) and using the “-m TEST” parameter to test for the best substitution model^40^ (Best-fit model: JTT+F+R5 chosen according to BIC). The tree was visualized using the iTOL program^41^. A similar analysis with sequences of 11 Rap genes from *B. subtilis* 168 used as an outgroup produced the same tree structure, and was used to define the tree root in Fig. 2a. Clades were manually defined for AloR receptors based on the branching patterns in the phylogenetic tree (Fig. 2a), as well as similarity in the cognate AloP sequence and the immediate genomic environment. For Fig. 2b, the dendrogram of the Alo receptors on the vertical axis of the matrix was generated by collapsing the tree shown in Fig. 2a. The dendrogram of the phylogenetic relationship between the *P. polymyxa* strains shown on the horizontal axis of Fig. 2b was generated by multiple sequence alignment and IQ-TREE analysis of the conserved GyrA protein of all 14 *P. polymyxa* strains, including GyrA from *B. subtilis* as an outgroup. For the data presented in Tables S1 and S3, AloR homologs were searched for their cognate *aloP* genes as described for *P. polymyxa* ATCC 842.

### Bacteria culture and growth conditions

*P. polymyxa* ATCC 842 was obtained from the BGSC (strain 25A2) and stored in −80°C as a glycerol stock. For all experiments, bacteria were first streaked on an LB plate, grown in 30°C at least overnight and colonies were used to inoculate round-bottom ventilated 15 mL tubes containing 4-5 mL LB medium, grown at 30°C with 200 RPM shaking.

### Preparation of growth media for mass spectrometry experiments

To search for peptide fragments secreted to the growth medium, overnight 5 mL cultures of *P. polymyxa* or *B. subtilis* 168 were first washed by centrifuging for 5 min in 3200 g at room temperature and re-suspended in 5 mL of chemically defined medium lacking any peptides described in reference ^64^. Washed cultures were then diluted 1:100 into 250 mL Erlenmeyer flasks with 50 mL defined media. The cells were grown in 30°C with 200 RPM shaking for 8 hrs for *P. polymyxa* log samples (OD ∼0.1) and 24 hrs for the stationary samples; and 5 hours for *B. subtilis* log samples (OD ∼0.3) and 8 hours for the stationary samples (OD ∼0.9). At the designated time point, 23 mL samples were centrifuged for 5 mins at 3200 g in 4°C, and two technical replicates of 10 mL of the supernatant for each sample were transferred to new tubes and flash frozen immediately.

### Sample preparation for MS-analysis

A method based on protein interaction with a solid phase was applied to enrich for peptides present in the supernatants. Prior to usage, a disposable solid phase enrichment (SPE) column (Phenomenex, 8B-S100-AAK), packed with an 8.5 nm pore size, modified styrene-divinylbenzene resin was equilibrated by rinsing with twice 1 mL acetonitrile and twice 1 mL water. Subsequently, 10 mL culture supernatant was loaded and unbound, and potentially larger proteins were removed by washing twice with 1 mL water. Finally, the enriched sample fraction was eluted with 0.5 mL 70% acetonitrile (v/v) and evaporated to dryness in a vacuum centrifuge. Samples were resuspended in 0.1% (v/v) acetic acid before MS analysis.

### Mass spectrometry

The enriched peptides were loaded on an EASY-nLC 1000 system (Thermo Fisher Scientific) equipped with an in-house built 20 cm column (inner diameter 100 mm, outer diameter 360 mm) filled with ReproSil-Pur 120 C18-AQ reversed-phase material (3 mm particles, Dr. Maisch GmbH, Germany). Elution of peptides was executed with a nonlinear 86 min gradient from 1 to 99% solvent B (0.1% (v/v) acetic acid in acetonitrile) with a flow rate of 300 nL/min and injected online into a QExactive mass spectrometer (Thermo Fisher Scientific). The survey scan at a resolution of R = 70,000 and 3 × 10^6^ automatic gain control target with activated lock mass correction was followed by selection of the 12 most abundant precursor ions for fragmentation. Data-dependent MS/MS scans were performed at a resolution of R = 17,500 and 1 × 10^5^ automatic gain control target with a NCE of 27.5. Singly charged ions as well as ions without detected charge states or charge states higher than six were excluded from MS/MS analysis. Dynamic exclusion for 30 s was activated.

### Database search of MS results

Identification of peptides was carried out by database search using MaxQuant 1.6.3.4 with the implemented Andromeda algorithm^65^ applying the following parameters: digestion mode unspecific; variable modification, methionine oxidation, and maximal number of 5 modifications per peptide; activated ‘match-between runs’ feature. The false discovery rates of peptide spectrum match, peptide and protein level were set to 0.01. Only unique peptides were used for identification. Two databases were used for peptide identification: a database for *P. polymyxa* ATCC 842 proteins downloaded from the IMG database^34^ and supplemented with the sequences for the Alo systems (5,429 entries); and the reference proteome of *B. subtilis* 168 downloaded from the Uniprot database (on 19/01/2019) containing 4,264 entries. Common laboratory contaminations and reverse entries were added during MaxQuant search and a peptide length of 5-35 amino acids was specified.

### Transcriptional response to peptides (RNA-seq experiments)

For RNA-seq experiments, synthetic lyophilized peptides were ordered from Peptide 2.0 Inc. (Chantilly, Virginia) and Genscript Corp at purity levels of 99% for the mature AloP13 (SHGRGG) and 81-93% for the AloP13 pro-peptide (SYFHNITIGNGSQITVSSHGRGG) and its scrambled version (VSGTRHGSFHSGIGNSGIYIQNT). The peptides were dissolved in DDW+2% DMSO to a working stock concentration of 100 µM and kept in aliquots in −20°C. Overnight *P. polymyxa* cultures grown in LB were diluted 1:100 into 500 mL Erlenmeyer flasks with 100 mL defined medium and grown in 30°C with 200 RPM shaking. At OD ∼0.1, 45 mL of each culture was washed by centrifuging for 5 min in 3,200 g at room temperature and re-suspending the pellet in 45 mL chemically defined medium^64^. The culture was split between 5 mL tubes containing each of the tested peptides at a final concentration of 5 µM, or a control tube containing a similar amount of DDW+2% DMSO. The cultures were incubated in 30°C with 200 RPM shaking for 10 min, after which 1:10 cold stop solution (90% (v/v) ethanol, 10% (v/v) saturated phenol) was added to the samples. The samples were centrifuged for 5 min at 3200 g in 4°C, the supernatant was discarded and the pellets were immediately flash frozen and stored in 80°C until RNA extraction. The experiment was conducted 3 times on different days to produce independent biological replicates.

Frozen bacterial pellets were lysed using the Fastprep homogenizer (MP Biomedicals, Santa Ana, California) and RNA was extracted with the FastRNA PRO(tm) blue kit (MP Biomedicals, 116025050) according to the manufacturer’s instructions. RNA levels and integrity were assessed using Qubit® RNA HS Assay Kit (Life technologies, Carlsbad, California, Q10210) and Tapestation (Agilent, Santa Clara, California, 5067-5576), respectively. All RNA samples were treated with TURBO(tm) DNase (Life technologies, AM2238). Ribosomal RNA depletion and RNA-seq libraries were prepared as described in ^66^, except that all reaction volumes were reduced by a factor of 4.

RNA-seq libraries were sequenced using Illumina NextSeq platform, and sequenced reads were demultiplexed using Illumina bcl2fastq module. Reads were mapped to the reference genome of *P. polymyxa* ATCC 842 downloaded from IMG (IMG taxon ID: 2547132099, 65 contigs), as described in ^66^. RNA-seq-mapped reads were used to generate genome-wide RNA-seq coverage maps and reads-per-gene counts.

Raw counts of reads-per-gene for each of the 3 biological replicates were used as input for DESeq2 package analysis using R 3.6.0^48^, while accounting for batch effect (DESeqDataSet (dds) design model “design= ∼batch + treatment”). Genes with normalized mean count <100 were discarded and a significant adjusted p-value (FDR<0.05) and fold change >2 or <0.5 were considered as the threshold for differentially expressed genes in each contrast (mature vs. control and scrambled vs. control). The volcano plot in Fig. 4a was plotted using EnhancedVolcano R package^67^, based on the results of the DESeqs2 analysis.

### Clumping assay

Cultures were incubated with peptides as described above for the RNA-seq experiments for 30 minutes, and 5 µL of culture were then dropped in the center of a dry LB plate and grown overnight in 30°C. The synthetic AloP13 pro-peptide lacking the last 6 aa (SYFHNITIGNGSQITVS) was ordered from Genscript Corp at a purity level of 76.6%. The plate was photographed using Biorad gel imager (Gel Doc XR+) and image colors were inverted for clarity.

## Supporting information

Table S1: AloR genes and corresponding AloP sequences identified in P. polymyxa strains

Table S2: AloR homologs in microbial genomes

Table S3: Alo systems in P. terrae NRRL B-30644

Table S4: All peptide fragments detected in mass spectrometry analysis

Table S5: Differential gene expression analysis at 10 minutes peptide incubation

Video S1: P. polymyxa clumping upon exposure to AloP13 pro-peptide

## Data Availability

All MS data (Fig. 3 and in Supplementary Table S4) have been deposited to the ProteomeXchange Consortium via the PRIDE^68^ partner repository with the dataset identifier PXD015319. All raw RNA-seq datasets (Fig. 4 and Supplementary Table S5) were deposited in the European Nucleotide Database (ENA), study accession no. PRJEB34369.

## Acknowledgements

We thank Yael Helman, Yaara Oppenheimer-Shaanan, Ohad Herches, Alon Savidor, Etai Rotem, Gilad Yaakov, Tabitha Bucher and Jürgen Bartel for their expertise and technical assistance. We thank the Sorek lab members for fruitful discussions and helpful suggestions. R.S. was supported, in part, by the Israel Science Foundation (personal grant 1360/16), the European Research Council (ERC) (grant ERC-CoG 681203), the German Research Council (DFG) priority program SPP 2002 (grant SO 1611/1-1), and the Knell Family Center for Microbiology. M.V. is a Clore Scholar and was supported by the Clore Israel Foundation. S.M. and D.B. were supported by the German Research Council (DFG) priority program SPP 2002 (grant BE 3869/5-1).

## Author Contributions

MV designed, conducted and analyzed all experiments unless otherwise indicated. SM and DB performed and analyzed the MS experiment, and TK ran the MS instrument. RS overviewed the project and was involved in the study design and data analyses. MV and RS wrote the paper.

## Supplementary Material

**Table S1**: AloR genes and corresponding AloP sequences identified in *P. polymyxa* strains

**Table S2**: AloR homologs in microbial genomes

**Table S3**: Alo systems in *P. terrae* NRRL B-30644

**Table S4**: All peptide fragments detected in mass spectrometry analysis

**Table S5:** Differential gene expression analysis at 10 minutes peptide incubation

**Video S1**: *P. polymyxa* clumping upon exposure to AloP13 pro-peptide

